# High pathogenicity avian influenza A (H5N1) clade 2.3.4.4b virus infection in a captive Tibetan black bear (*Ursus thibetanus*): investigations based on paraffin-embedded tissues, France, 2022

**DOI:** 10.1101/2023.10.19.563114

**Authors:** Pierre Bessière, Nicolas Gaide, Guillaume Croville, Manuela Crispo, Maxime Fusade-Boyer, Yanad Abou Monsef, Malorie Dirat, Marielle Beltrame, Philippine Dendauw, Karin Lemberger, Jean-Luc Guérin, Guillaume Le Loc’h

## Abstract

High pathogenicity avian influenza viruses H5Nx (HPAIVs) of clade 2.3.4.4b have been circulating increasingly in both wild and domestic birds in recent years. In turn, this has led to an increase in the number of spillovers events affecting mammals. In November 2022, a HPAIV H5N1 caused an outbreak in a zoological park in the south of France, resulting in the death of a Tibetan black bear (*Ursus thibetanus*) and several captive and wild bird species. We detected the virus in various tissues of the bear and a wild black-headed gull found dead in its enclosure using histopathology, two different *in-situ* detection techniques and next generation sequencing, all performed on formalin fixed paraffin embedded tissues. Phylogenetic analysis performed on HA gene segment showed that bear and gull strains shared 99.998% genetic identity, making the bird strain the closest related one. We detected the PB2 E627K mutation in minute quantities in the gull, whereas it predominated in the bear, which suggests that this mammalian adaptation marker was selected during the bear infection. Our results provide the first molecular and histopathological characterization of an H5N1 virus infection in this bear species.

## Introduction

The genetic and antigenic diversity of influenza viruses is considerable: 16 and 9 hemagglutinin and neuraminidase subtypes respectively circulate in wild waterfowl, considered the reservoir of influenza viruses [1]. Following the acquisition of a mutation in the sequence encoding the hemagglutinin cleavage site, viruses belonging to the H5Nx and H7Nx subtypes are capable of acquiring a high pathogenicity phenotype. While the tropism of low pathogenicity avian influenza viruses (LPAIVs) is mainly restricted to the digestive and respiratory tracts, high pathogenicity avian influenza viruses (HPAIVs) can replicate systemically [2].

For a long time, HPAIVs circulating in domestic birds were considered unlikely to return to the wild compartment: viruses adapt to the species they infect, and many viruses adapted to Gallinaceae are poorly adapted to wild waterfowl [3]. However, in recent years, this dogma has been shaken: HPAI H5Nx viruses, which initially appeared in the domestic compartment, have succeeded in becoming endemic in wild birds [4]. More importantly, while the species barrier between mammals and birds is considerable, these viruses have managed to cross it on several occasions.

H5Nx HPAIVs’ circulation, particularly those of clade 2.3.4.4b, has significantly increased in recent years: viruses of this clade are spreading in wild bird populations more rapidly than ever since their emergence in 1996 [5]. Thus, the occasions on which mammals have been exposed to the virus have become more frequent in turn. Infection generally occurs following exposure to contaminated feces or water, or animal carcasses [6]. Sporadic infections of wild mammals have been described in the past, but have never been as frequent as in recent years. Numerous domestic and wild animals have been contaminated, including various species of seals, sea lions, foxes, cats, raccoons, skunks and bears [7– 12].

These sporadic infections are not always dead ends [13], and every time an avian influenza virus manages to infect a mammal, there is a risk that adaptive mutations will appear [1]. When enough genetic changes accumulate, the result can be the emergence of a virus more efficiently transmitted between mammals. Ultimately, a novel influenza virus able to sustain transmission between humans could cause a pandemic [14].

In November 2022, a Tibetan black bear (*Ursus thibetanus*) from the Sigean zoo in France was reported dead. Although infection with an avian influenza virus was not initially suspected, post-mortem analyses revealed the presence of clade 2.3.4.4b HPAIV H5N1 in various organs and blood. Over the following fortnight, several influenza-positive birds were also found dead nearby the bear enclosure suggesting a bird-to-bear transmission. Using formalin fixed paraffin embedded tissue, we conducted molecular and histopathological investigation to characterize this outbreak.

## Results

### Outbreak detection

In early November 2022, a 12-year-old male Tibetan black bear (*Ursus thibetanus*) was found dead in its enclosure at the Sigean zoo, France. A few days before, zookeepers had noticed a slight decline in general condition, while the day prior to its death, the bear presented marked dyspnea, hyperthermia, lateral decubitus and diarrhea. Analyses carried out by the zoo’s veterinarians showed severe leukopenia and hypercreatininemia, consistent with acute renal failure (**Supplementary File 1**). A necropsy was carried out, revealing hemorrhagic lesions (petechiae and suffusion) on the epicardium and liver, severe congestion of lungs, kidneys and intestines, and finally, a necrotic tracheal mucosa. Over the following days, several zoo and wild birds (pelicans, jackdaws and gulls) died and tested positive by RT-qPCR for clade 2.3.4.4b HPAIV H5N1, one of the carcasses being found in the bear’s enclosure: a black-headed gull (*Chroicocephalus ridibundus*), which the zoo veterinarians also necropsied (mark ed pulmonary and splenic congestion were the most remarkable findings identified at necropsy). In the days following the bear’s death, other bears of the same species displayed mild to moderate clinical respiratory signs, but could not be sampled to be analyzed as part of this study.

The possibility that the bear might have been infected by an H5N1 virus of clade 2.3.4.4.b was subsequently suspected, and confirmed by molecular analysis of biological samples sent to the French reference laboratory for high pathogenicity avian influenza, which deposited the viral genome sequence on GISAID (isolate ID EPI_ISL_17233426). After initial routine screening at a diagnostic histopathology laboratory (Vet Diagnostics, France), formalin fixed paraffin-embedded (FFPE) tissues from the bear and gull were then sent to the ENVT laboratory for further investigation.

### Pathological examination of the infected bear and black-headed gull

Histopathological examination of the bear revealed acute, marked multifocal to coalescing fibrino-necrotizing lymphadenitis and splenitis, associated with vasculitis and hemorrhages (**Figure 1)**; marked pulmonary edema and congestion; moderate suppurative tracheitis with concurrent submucosal vascular thrombosis; severe, multifocal necro-suppurative hepatitis. Marked acute hemorrhages, involving the subepicardium and renal interstitium, were also present, while the gastro-intestinal tract appeared unremarkable (**Supplementary Figure 1**). Viral antigen and RNA were frequently detected in a visceral lymph node, multifocally, within areas of necrotizing vasculitis (**Figure 1**). In the lung, viral antigen was sparsely detected within bronchoalveolar luminal debris, lobular and interlobular interstitium admixed with non-specific background. Viral RNA was similarly detected in terms of distribution, although the signal appeared more widespread within the pulmonary parenchyma (**Figure 1**). In the kidney, viral antigen was rarely observed within a few glomerular tufts, while viral RNA detection was negative (**Supplementary Figure 1**). Additionally, RNAscope *in situ* hybridization revealed sparse viral RNA within interstitium of myocardium, gastro-intestinal and tracheal mucosa and submucosa and splenic red pulp. Other findings included myocardial atherosclerosis and mineralization.

**Figure 1.**
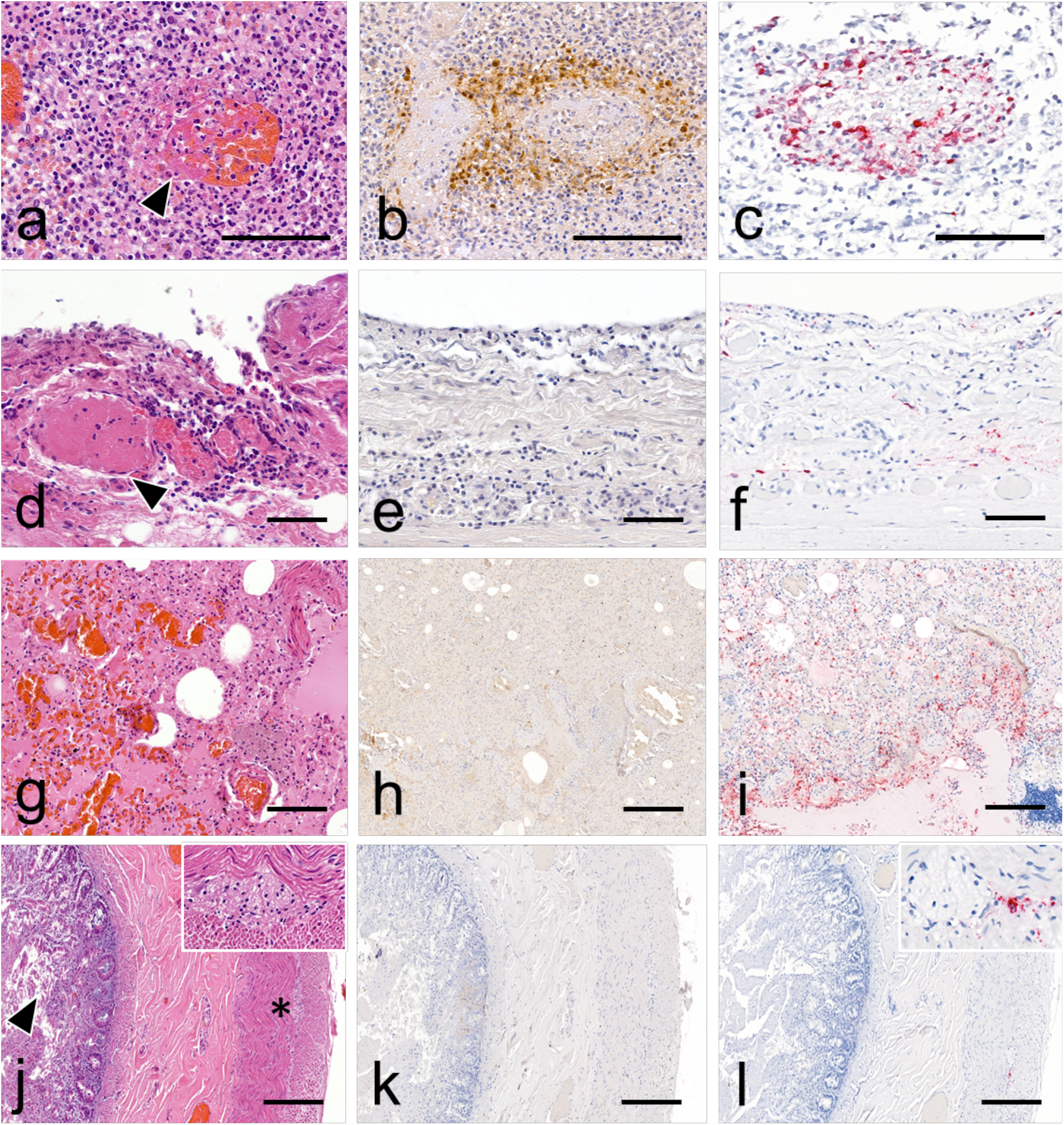
Histopathology, viral antigen and RNA detection in tissues obtained from infected bear. a. Visceral lymph node: necrotizing vasculitis (arrowhead) with thrombosis and hemorrhages. Hematoxylin and eosin (H&E) stain. b. Visceral lymph node: viral antigen is observed associated with vasculitis and extending to the surrounding lympho-nodal parenchyma. Anti-nucleoprotein influenza A immunohistochemistry (anti-NP IHC). c. Visceral lymph node: viral RNA is intralesionally detected within areas of vasculitis. M gene RNAscope *in situ* hybridization (RNAscope ISH). d. trachea: thrombosis (arrowhead) and perivascular leukocytic infiltration are observed within the mucosa and submucosa. The overlying epithelium is sloughed (arrowhead) (H&E stain). e. Trachea: no viral antigen detection is observed (anti-NP IHC). f. Trachea: positive viral RNA detection is observed in the interstitium of mucosa and submucosa (RNAscope ISH). g. Lung: diffuse congestion and edema (H&E stain). h. Lung: IHC shows moderate non-specific background staining with no significant detection of viral antigen at low magnification (anti-NP IHC). i. Lung: viral RNA is widely distributed within the lobular and interlobular interstitium (RNAscope ISH). Intestine: autolytic changes are present in the mucosa, including cell sloughing (arrowhead). The submucosa, tunica muscularis and serosa appear within normal limits. The myenteric plexus (insert and asterisk) is readily identifiable and also normal (H&E stain). j. Intestine: no viral antigen detection is observed (anti-NP IHC). l. Intestine: Viral RNA is focally present within the myenteric plexus (insert) (RNAscope ISH). Scale bars: 50μm (d-f), 100 μm (a-c, g-i), 200 μm (j-l).

Histopathological assessment of the gull showed acute necrotic-inflammatory changes in the majority of the organs examined. Mild to marked encephalitis, pancreatitis, splenitis, hepatitis, nephritis and thyroiditis lesions exhibited variable amounts of viral antigen and RNA highlighted by IHC and RNAscope ISH, respectively (**Supplementary Figures 2 and 3**).

**Figure 2:**
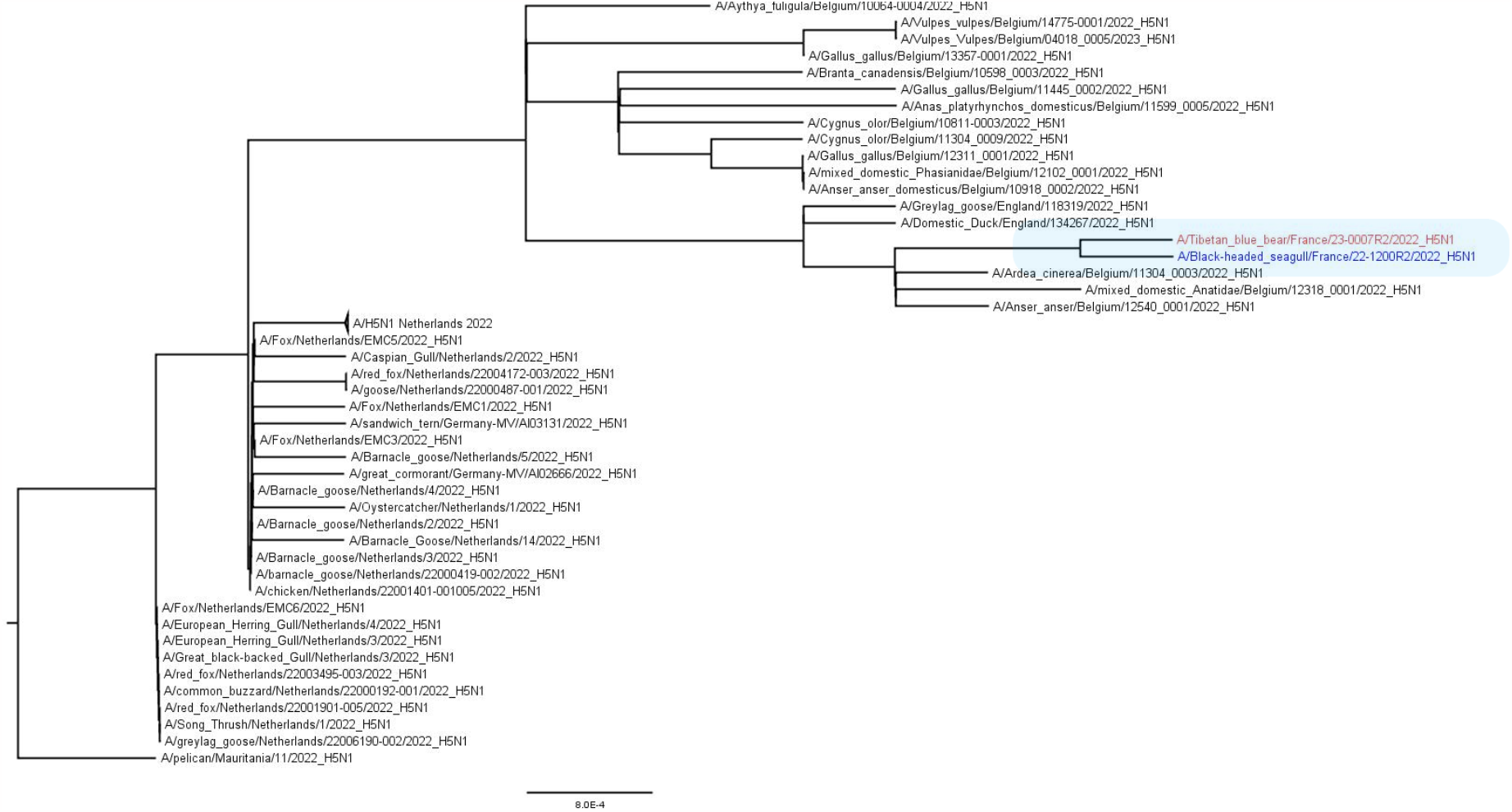
HA maximum likelihood phylogenetic tree. Bear and gull-derived sequences are labelled in red and blue respectively. Scale bar: number of nucleotide substitution per site.

### Phylogenetic and genetic analyses

We performed next generation sequencing on several bear samples (lymph node, lung, and liver) and one gull sample (brain), using an Element AVITI sequencer (Element Biosciences, San Diego, CA) and a 2x150 bp paired-end protocol. We found H5N1 virus reads in all samples, in sufficient numbers to reconstitute whole genome consensus sequences, available under accession numbers OR634756 to OR634763 and OR634764 to OR634771 for the bear and the gull isolates, respectively. Details of the viral (>100 reads) and bacterial (>1000 reads) species identified by metagenomics can be found in **Supplementary File 2**.

Phylogenetic analysis performed on HA gene segment showed that viruses found on both animals belonged to clade 2.3.4.4b. The HA sequence of the bear-derived virus shared 99.998% genetic identity with the HA of the strain detected in the gull, making the bird strain the closest related strain (**Figure 2**). This finding was further confirmed by the phylogenetic analysis performed on the other viral segments, which showed that both strains were closely related to strains circulating in Belgium at that time (**Supplementary Figure 5**). The whole genome analysis revealed both strains belonged to the AB genotype (H5N1-A/duck/Saratov/29-02/2021-like), the main circulating genotype at that time in Europe [6].

Analysis of the consensus sequences revealed the presence of several mammalian adaptation markers, listed in **Table 1**, in both bear and gull viruses, markers also found in the genomes of other phylogenetically related avian viruses. However, only the bear virus possessed the PB2 E627K marker, suggesting that this mutation had emerged over the course of infection.

**Table 1:** Mammalian adaptation markers found on the gull and bear derived viral sequences. HA sequence numbering was performed including the peptide signal.

| Protein | Amino-acid position | Gull | Bear | Reference |
| --- | --- | --- | --- | --- |
| PB2 | 627 | E | K | [15] |
| PB1-F2 | 66 | S | S | [16] |
| HA | 149 | A | A | [17] |
|  | 170 | N | N | [18] |
|  | 172 | A | A |  |
| M1 | 30 | D | D | [19] |
|  | 215 | A | A |  |
| NS1 | 42 | S | S | [20] |
|  | 92 | D | D | [21] |
|  | 103 | F | F | [22] |
|  | 106 | M | M |  |

We then looked for the presence of minority variants using variant calling analysis. While this analysis did not reveal any other mammalian adaptation markers already described in the literature, it did show the presence of variants with either an E or a K at position 627 of the PB2 protein, in both bear and gull viruses (**Supplementary File 3**). More specifically, in the gull, over 99% of reads coded for an E, while in the bear, 37.3% coded for an E and 62.7% coded for a K at position 627, both in the liver and lung samples. Although the sequencing depth at this position was around 1400 reads, no sequence encoding an E at position 627 was found in the lymph node sample.

## Discussion

We revealed the presence of clade 2.3.4.4b H5N1 virus in various tissues of a Tibetan black bear and a black-headed gull, using histopathology, two different *in-situ* detection techniques and next generation sequencing. Sequence analysis revealed that the viruses found in the bear and gull were phylogenetically very close, and that variants with the PB2 E627K mutation had emerged in the bear. Our findings do not allow us to determine with any certainty how the bear became infected, especially as the gull was found dead several days after the bear. The most plausible hypothesis is that the virus circulated undetected for several days in the zoo’s wild and captive avifauna, and that the bear was either in direct contact with an infected bird (dead or alive), or in contact with water or food contaminated by avian droppings. Interestingly, the PB2 E627K mutation was present in very small amounts in the gull sample, whereas it was predominant in the bear lung sample. We therefore believe that it did not arise in the bear through *de novo* mutations, but that the bear’s infection was seeded by an inoculum containing PB2 627E variants and a few PB2 627K variants, which were then selected because of their selective advantage [15]. Interestingly, the bear lymph node sample contained no PB2 627E variants. This suggests that the PB2 627K variant was selected in the respiratory tract, from which it then spread systemically.

When an animal is necropsied outside a research facility, improper preservation of samples is a frequent issue, especially if the carcasses cannot be refrigerated and if the analysis cannot be performed quickly. One of the advantages of formalin-fixed paraffin-embedded specimens is that they can be stored at room temperature for years, if not decades, and are shipped easily [27]. Although this process degrades the nucleic acids to some extent, methods for extracting DNA and RNA of sufficient quality are now available [28]. This is how we managed to use FFPE tissues to reconstruct an H5N1 avian influenza virus outbreak in a zoological park through molecular investigations, despite the absence of fresh tissue samples.

Another limitation of our study is that the bear’s brain was not examined and sampled. Collecting this organ would have been technically challenging and since the zoo’s veterinary service did not suspect an infection with an avian influenza virus, at the time of the necropsy, the cranium was not opened. However, the involvement of the central nervous system has been frequently identified in both birds and mammals infected with an H5Nx virus of clade 2.3.4.4b, with neurological disorders sometimes being the only clinical signs observed [28]. For this reason, highlight the presence of the virus and concurrent histopathological lesions in the brain of our bear would have been of particular interest.

In zoological parks many species may reside in close proximity to one another and avoiding contact with wild birds is difficult, if not impossible, when animals are not kept in cages, but in open spaces. Zookeepers should receive proper training in biosecurity and made aware of the threats posed by avian influenza viruses toward mammals, as they were towards SARS-CoV-2 during the COVID-19 pandemic [29]. This aspect is important, not only for protecting animals, particularly in case of endangered species, but also for preventing epizootic flare-ups. As the RNA polymerase of influenza viruses is prone to errors during viral genome replication, better adapted variants may appear when a mammal is infected by an avian influenza virus [30]. When such a virus spreads from mammal to mammal, the risk of a more transmissible variant being selected increases considerably, and chains of transmission must be avoided as much as possible [31]. By raising awareness among veterinarians and zookeepers of the clinical presentations associated with H5Nx virus infection in mammals, the number of undetected epizootics should be reduced, with zoological parks thus acting as sentinels.

In conclusion, biosecurity and surveillance programs are essential to deal with epizootics caused by clade 2.3.4.4b H5Nx viruses, and the zoonotic spillovers that are becoming increasingly frequent. In particular, active and passive surveillance of mammals, including both wild, captive and domestic, will be invaluable in anticipating the emergence of H5Nx viruses with pandemic potential [33].

## Materials and methods

### Histopathology

Tissue samples were placed in 10% neutral buffered formalin. For the bear, available tissues included trachea, lung, heart, visceral lymph node, spleen, intestine, stomach and kidney. For the gull brain, trachea, lung, heart, spleen, pancreas (splenic lobe), thyroid gland, liver, intestine and kidney were collected. After fixation, tissues were routinely processed in paraffin blocks, sectioned at 3μm, stained with hematoxylin and eosin (H&E) and examined by light microscopy.

### Immunohistochemistry

To assess viral antigen tissue distribution within the bear and gull tissues, immunohistochemistry (IHC) was performed on formalin-fixed paraffin-embedded (FFPE) tissue sections, using a monoclonal mouse anti-nucleoprotein influenza A virus antibody (Biozol BE0159, pronase 0.05% retrieval solution, 10 min at 37°C: antibody dilution 1/2000, incubation overnight at 4°C). The immunohistochemical staining was revealed with HRP labeled polymer (EnVisio+ Dual Link System HRP, K4061, Agilent) and the diaminobenzidine HRP chromogen (DAB+ liquid, K3467, Agilent). Negative controls included sections incubated either without the primary antibody or with another monoclonal antibody of the same isotype (IgG2).

### RNAscope ISH

To determine the presence of Avian Influenza A Virus RNA and assess its distribution within the bear tissue sections, RNAscope *in situ* hybridization (RNAscope ISH) was performed as previously described [23]. Briefly, we used probes targeting M1 and M2 genes (V-InfluenzaA-H5N8-M2M1 probe), H5 hemagglutinin gene (V-InfluenzaA-H5N8-HA-O1 probe) of clade 2.3.4.4b HPAIV H5 and a RNAscope 2.5 high-definition red assay, according to the manufacturer’s instructions, including mild pretreatment conditions (15 min incubation with protease digestion for antigenic retrieval) and hematoxylin counterstaining. A probe targeting the dihydrodipicolinate reductase (dapB) gene from the *Bacillus subtilis* strain SMY, served as negative control.

### Next generation sequencing

Three bear samples and one gull sample were selected for the metagenomics analysis on the RNA fraction: lymph node (bear), lung (bear), liver (bear) and brain (gull). One bear sample (liver) was selected for the metagenomics analysis on the DNA fraction.

The nucleic acids were extracted from the FFPE tissue sections using the Nucleospin total RNA FFPE XS (Macherey-Nagel). RNA sequencing libraries were prepared using the NEBNext® Ultra™ II Directional RNA Library Prep Kit (New England Biolabs) and the DNA library was prepared using the NEBNext® Microbiome DNA Enrichment Kit (New England Biolabs). The sequencing run was then performed on the Element AVITI sequencer (Element Biosciences, San Diego, CA) using a 2x150 bp paired-end protocol.

### Bioinformatics analysis

The metagenomic data analysis was performed with Kraken2 [24]. We set a threshold at 100 and 1000 reads for the abundance of the viral and bacterial species, respectively. The reads were then mapped on a H5N1 reference genome (GISAID isolate ID: EPI_ISL_17233426) with minimap2 [25] and the consensus sequences were generated using iVar [26].

### Phylogenetic analysis

Consensus sequences of each viral gene segment detected in black bear were compared with the most related sequences available in GISAID (https://www.gisaid.org/) and aligned by using MAFFT version 7 (https://mafft.cbrc.jp/alignment/server/index.html). Maximum-likelihood phylogenetic trees were generated using RaxML, version 8.2.X (https://sco.h-its.org/exelixis/software.html), with a GTR model associated with gamma distribution (https://github.com/tamuri/treesub/blob/master/README.md. Phylogenetic trees were then visualized by using FigTree version 1.4.2.

## Acknowledgment

This study was performed in the framework of the “Chaire de Biosécurité et Santé Aviaires”, hosted by the National Veterinary College of Toulouse (ENVT) and funded by the Direction Générale de l’Alimentation, Ministère de l’Agriculture et de la Souveraineté Alimentaire, France.

The authors would like to thank HELIXIO SAS (Saint Beauzire—France) for performing the metagenomics analysis.

## Author contributions

P.B., N.G., G.C., M.C., J.L.G. and G.L.L. conceptualized and designed experiments. P.B., N.G., G.C., M.C.,

M.F.B., M.D., K.L., M.B., P.D. performed experiments. P.B., N.G., M.C., Y.A.M., G.C., M.F.B., K.L., J.L.G. and G.L.L. analyzed data. J.L.G. and G.L.L. acquired funding. P.B., N.G., M.C. and G.C. drafted the manuscript. All authors reviewed and edited the final manuscript before submission.

