## Supplementary File 1 for "High pathogenicity avian influenza A (H5N1) clade 2.3.4.4b virus infection in a captive Tibetan black bear (*Ursus thibetanus*): investigations based on paraffin-embedded tissues, France, 2022"

**Complete blood count:**
Red blood cells: 8.01 x $10^{12}$/L
Hematocrit: 4.8%
Hemoglobin: 17.6g/dL
Mean corpuscular volume: 59.7fL
Mean corpuscular hemoglobin: 21.9pg
Mean corpuscular hemoglobin concentration: 36.7g/dL
Red cell distribution width: 16.1%
% Reticulocyte: 16.1%
Reticulocytes: 31,6 K/µL
White blood cells: 1.80 x $10^{9}$/L
- Neutrophiles: 37.2%
- Lymphocytes: 56.0%
- Monocytes: 2.9%
- Eosinophiles: 3.9%
- Basophiles: 0%

Platelets: 36 K/µL

**Blood biochemistry:**
Glucose: 1.52g/L
Creatinine: 60mg/L
Urea: 0.436g/L
Total protein: 83g/L
Albumin: 39g/L
Globulin: 44g/L
Alanine transaminase: 225U/L
Alkaline phosphatase: 41U/L
C reactive protein: < 1mg/L
