## Supplementary Figure 4 for "High pathogenicity avian influenza A (H5N1) clade 2.3.4.4b virus infection in a captive Tibetan black bear (*Ursus thibetanus*): investigations based on paraffin-embedded tissues, France, 2022"

Maximum likelihood phylogenetic trees performed on viral segments other than HA. Bear and gull-derived sequences are labelled in red and blue respectively. Scale bar: number of nucleotide substitution per site.

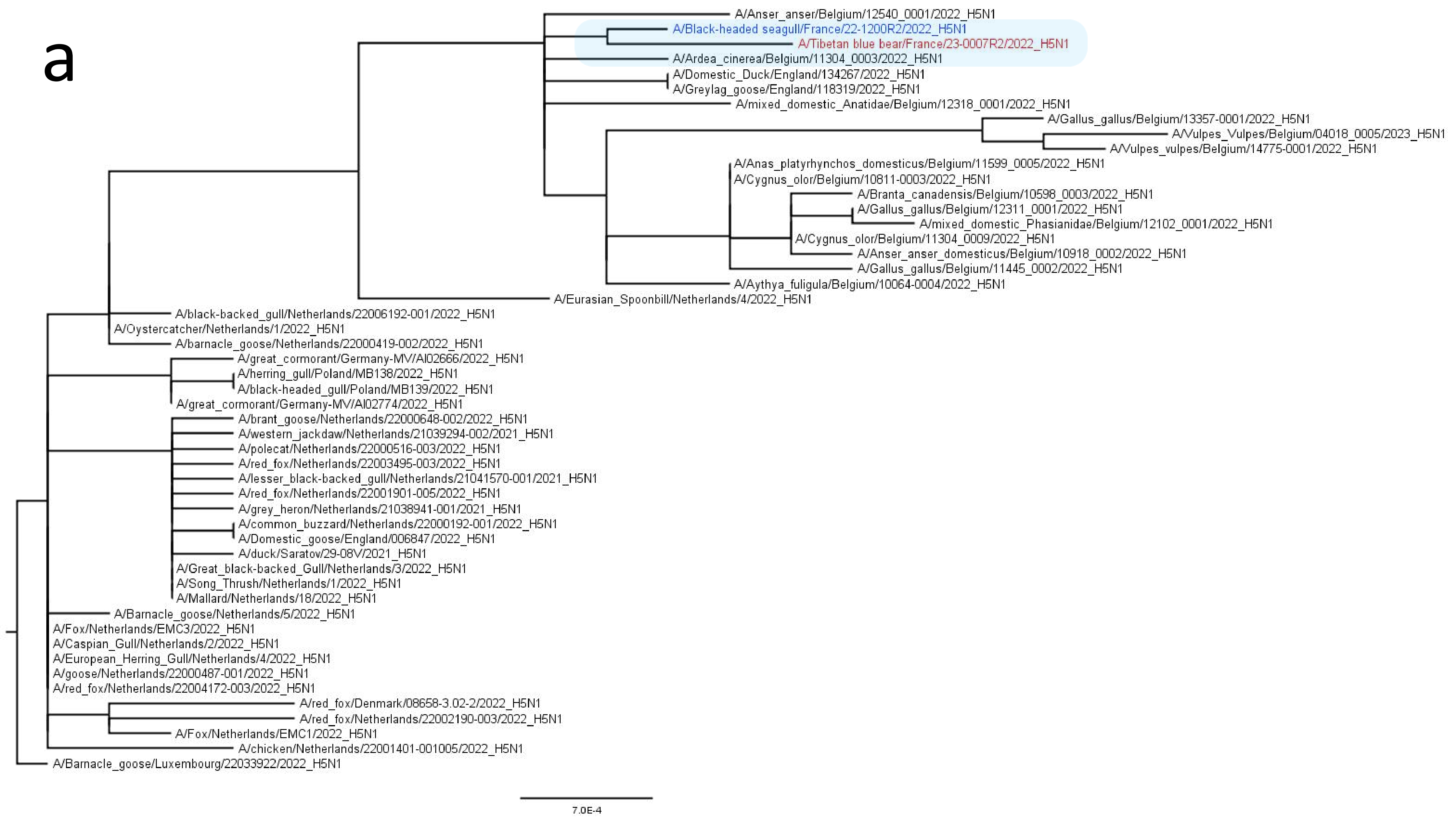

a
A/Anser_anser/Belgium/12540_0001/2022_H5N1
A/Black-headed seagull/France/22-1200R2/2022_H5N1
A/Tibetan blue bear/France/23-0007R2/2022_H5N1
A/Ardea_cinerea/Belgium/11304_0003/2022_H5N1
A/Domestic_Duck/England/134267/2022_H5N1
A/Greylag_goose/England/118319/2022_H5N1
A/mixed_domestic_Anatidae/Belgium/12318_0001/2022_H5N1
A/Gallus_gallus/Belgium/13357-0001/2022_H5N1
A/Vulpes_Vulpes/Belgium/04018_0005/2023_H5N1
A/Vulpes_vulpes/Belgium/14775-0001/2022_H5N1
A/Anas_platyrhynchos_domesticus/Belgium/11599_0005/2022_H5N1
A/Cygnus_olor/Belgium/10811-0003/2022_H5N1
A/Branta_canadensis/Belgium/10598_0003/2022_H5N1
A/Gallus_gallus/Belgium/12311_0001/2022_H5N1
A/mixed_domestic_Phasianidae/Belgium/12102_0001/2022_H5N1
A/Cygnus_olor/Belgium/11304_0009/2022_H5N1
A/Anser_anser_domesticus/Belgium/10918_0002/2022_H5N1
A/Gallus_gallus/Belgium/11445_0002/2022_H5N1
A/Aythya_fuligula/Belgium/10064-0004/2022_H5N1
A/Eurasian_Spoonbill/Netherlands/4/2022_H5N1
A/black-backed_gull/Netherlands/22006192-001/2022_H5N1
A/Oystercatcher/Netherlands/1/2022_H5N1
A/barnacle_goose/Netherlands/22000419-002/2022_H5N1
A/great_cormorant/Germany-MV/AI02666/2022_H5N1
A/herring_gull/Poland/MB138/2022_H5N1
A/black-headed_gull/Poland/MB139/2022_H5N1
A/great_cormorant/Germany-MV/AI02774/2022_H5N1
A/brant_goose/Netherlands/22000648-002/2022_H5N1
A/western_jackdaw/Netherlands/21039294-002/2021_H5N1
A/polecat/Netherlands/22000516-003/2022_H5N1
A/red_fox/Netherlands/22003495-003/2022_H5N1
A/lesser_black-backed_gull/Netherlands/21041570-001/2021_H5N1
A/red_fox/Netherlands/22001901-005/2022_H5N1
A/grey_heron/Netherlands/21038941-001/2021_H5N1
A/common_buzzard/Netherlands/22000192-001/2022_H5N1
A/Domestic_goose/England/006847/2022_H5N1
A/duck/Saratov/29-08V/2021_H5N1
A/Great_black-backed_Gull/Netherlands/3/2022_H5N1
A/Song_Thrush/Netherlands/1/2022_H5N1
A/Mallard/Netherlands/18/2022_H5N1
A/Barnacle_goose/Netherlands/5/2022_H5N1
A/Fox/Netherlands/EMC3/2022_H5N1
A/Caspian_Gull/Netherlands/2/2022_H5N1
A/European_Herring_Gull/Netherlands/4/2022_H5N1
A/goose/Netherlands/22000487-001/2022_H5N1
A/red_fox/Netherlands/22004172-003/2022_H5N1
A/red_fox/Denmark/08658-3.02-2/2022_H5N1
A/red_fox/Netherlands/22002190-003/2022_H5N1
A/Fox/Netherlands/EMC1/2022_H5N1
A/chicken/Netherlands/22001401-001005/2022_H5N1
A/Barnacle_goose/Luxembourg/22033922/2022_H5N1
7.0E-4

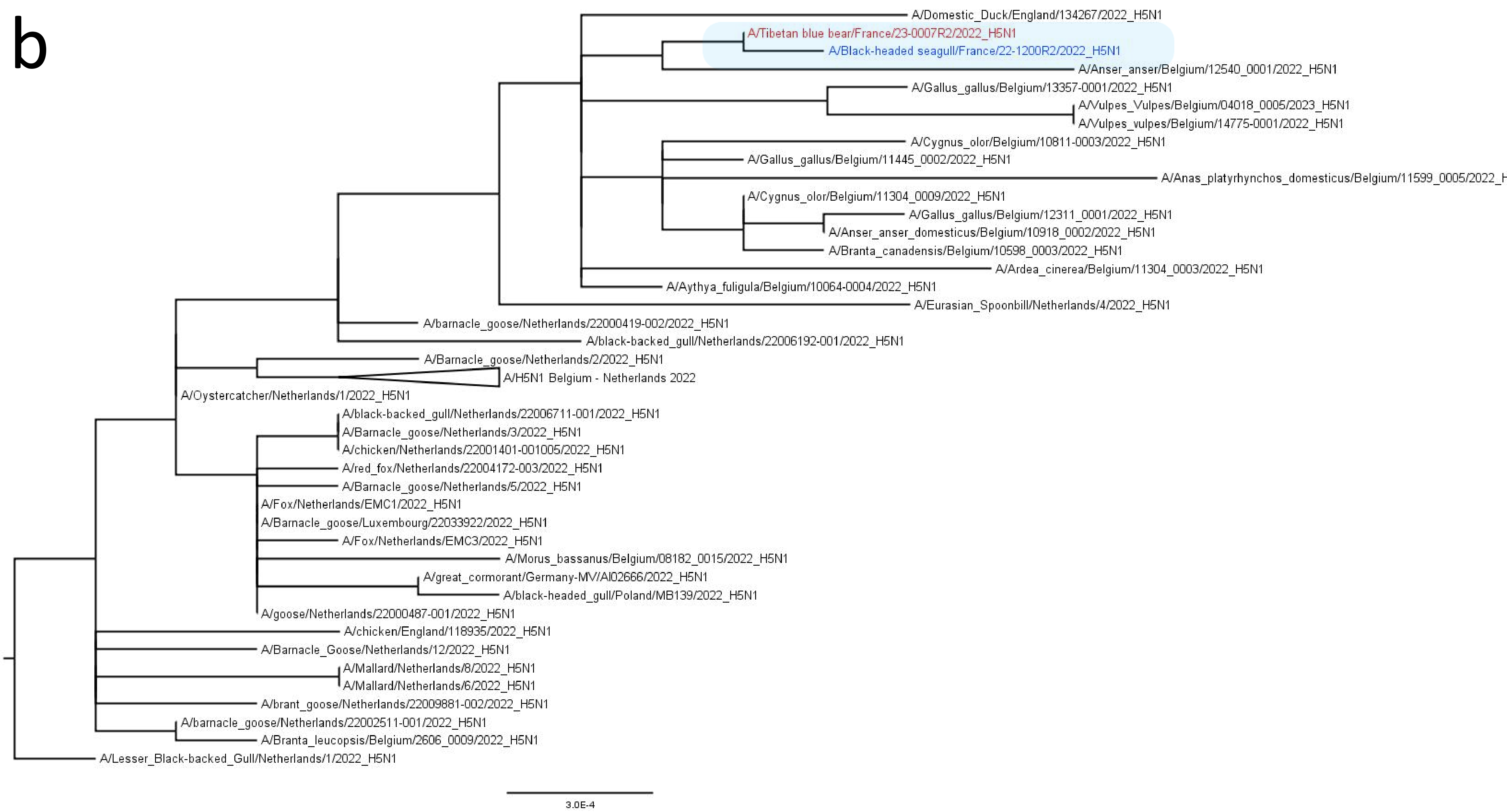
b
A/Domestic_Duck/England/134267/2022_H5N1
A/Tibetan blue bear/France/23-0007R2/2022_H5N1
A/Black-headed seagull/France/22-1200R2/2022_H5N1
A/Anser_anser/Belgium/12540_0001/2022_H5N1
A/Gallus_gallus/Belgium/13357-0001/2022_H5N1
A/Vulpes_Vulpes/Belgium/04018_0005/2023_H5N1
A/Vulpes_vulpes/Belgium/14775-0001/2022_H5N1
A/Cygnus_olor/Belgium/10811-0003/2022_H5N1
A/Gallus_gallus/Belgium/11445_0002/2022_H5N1
A/Anas_platyrhynchos_domesticus/Belgium/11599_0005/2022_H
A/Cygnus_olor/Belgium/11304_0009/2022_H5N1
A/Gallus_gallus/Belgium/12311_0001/2022_H5N1
A/Anser_anser_domesticus/Belgium/10918_0002/2022_H5N1
A/Branta_canadensis/Belgium/10598_0003/2022_H5N1
A/Ardea_cinerea/Belgium/11304_0003/2022_H5N1
A/Aythya_fuligula/Belgium/10064-0004/2022_H5N1
A/Eurasian_Spoonbill/Netherlands/4/2022_H5N1
A/barnacle_goose/Netherlands/22000419-002/2022_H5N1
A/black-backed_gull/Netherlands/22006192-001/2022_H5N1
A/Barnacle_goose/Netherlands/2/2022_H5N1
A/H5N1 Belgium - Netherlands 2022
A/Oystercatcher/Netherlands/1/2022_H5N1
A/black-backed_gull/Netherlands/22006711-001/2022_H5N1
A/Barnacle_goose/Netherlands/3/2022_H5N1
A/chicken/Netherlands/22001401-001005/2022_H5N1
A/red_fox/Netherlands/22004172-003/2022_H5N1
A/Barnacle_goose/Netherlands/5/2022_H5N1
A/Fox/Netherlands/EMC1/2022_H5N1
A/Barnacle_goose/Luxembourg/22033922/2022_H5N1
A/Fox/Netherlands/EMC3/2022_H5N1
A/Morus_bassanus/Belgium/08182_0015/2022_H5N1
A/great_cormorant/Germany-MV/AI02666/2022_H5N1
A/black-headed_gull/Poland/MB139/2022_H5N1
A/goose/Netherlands/22000487-001/2022_H5N1
A/chicken/England/118935/2022_H5N1
A/Barnacle_Goose/Netherlands/12/2022_H5N1
A/Mallard/Netherlands/8/2022_H5N1
A/Mallard/Netherlands/6/2022_H5N1
A/brant_goose/Netherlands/22009881-002/2022_H5N1
A/barnacle_goose/Netherlands/22002511-001/2022_H5N1
A/Branta_leucopsis/Belgium/2606_0009/2022_H5N1
A/Lesser_Black-backed_Gull/Netherlands/1/2022_H5N1
3.0E-4

C

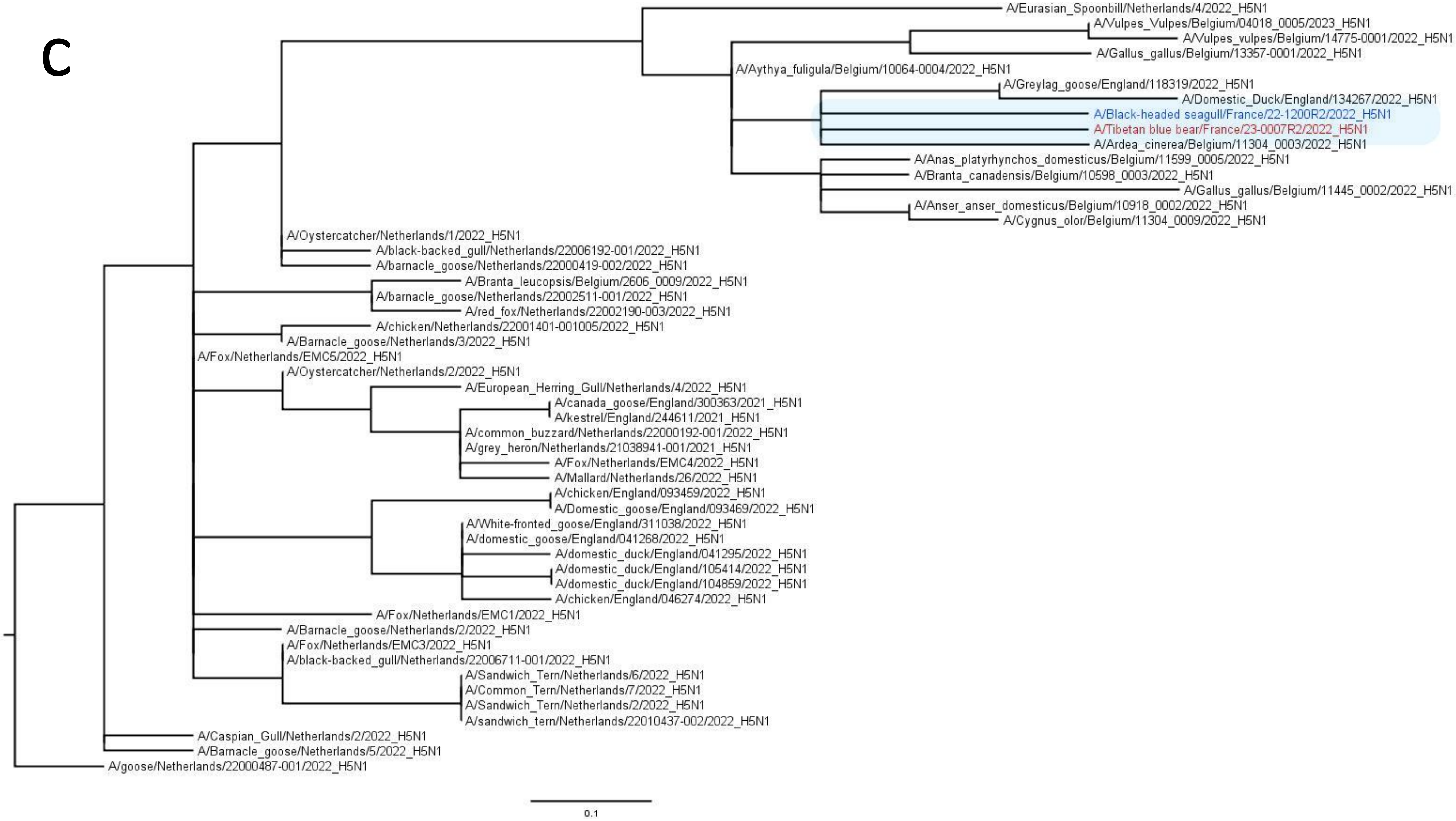
A/Eurasian_Spoonbill/Netherlands/4/2022_H5N1
A/Vulpes_Vulpes/Belgium/04018_0005/2023_H5N1
A/Vulpes_vulpes/Belgium/14775-0001/2022_H5N1
A/Gallus_gallus/Belgium/13357-0001/2022_H5N1
A/Aythya_fuligula/Belgium/10064-0004/2022_H5N1
A/Greylag_goose/England/118319/2022_H5N1
A/Domestic_Duck/England/134267/2022_H5N1
A/Black-headed seagull/France/22-1200R2/2022_H5N1
A/Tibetan blue bear/France/23-0007R2/2022_H5N1
A/Ardea_cinerea/Belgium/11304_0003/2022_H5N1
A/Anas_platyrhynchos_domesticus/Belgium/11599_0005/2022_H5N1
A/Branta_canadensis/Belgium/10598_0003/2022_H5N1
A/Gallus_gallus/Belgium/11445_0002/2022_H5N1
A/Anser_anser_domesticus/Belgium/10918_0002/2022_H5N1
A/Cygnus_olor/Belgium/11304_0009/2022_H5N1
A/Oystercatcher/Netherlands/1/2022_H5N1
A/black-backed_gull/Netherlands/22006192-001/2022_H5N1
A/barnacle_goose/Netherlands/22000419-002/2022_H5N1
A/Branta_leucopsis/Belgium/2606_0009/2022_H5N1
A/barnacle_goose/Netherlands/22002511-001/2022_H5N1
A/red_fox/Netherlands/22002190-003/2022_H5N1
A/chicken/Netherlands/22001401-001005/2022_H5N1
A/Barnacle_goose/Netherlands/3/2022_H5N1
A/Fox/Netherlands/EMC5/2022_H5N1
A/Oystercatcher/Netherlands/2/2022_H5N1
A/European_Herring_Gull/Netherlands/4/2022_H5N1
A/canada_goose/England/300363/2021_H5N1
A/kestrel/England/244611/2021_H5N1
A/common_buzzard/Netherlands/22000192-001/2022_H5N1
A/grey_heron/Netherlands/21038941-001/2021_H5N1
A/Fox/Netherlands/EMC4/2022_H5N1
A/Mallard/Netherlands/26/2022_H5N1
A/chicken/England/093459/2022_H5N1
A/Domestic_goose/England/093469/2022_H5N1
A/White-fronted_goose/England/311038/2022_H5N1
A/domestic_goose/England/041268/2022_H5N1
A/domestic_duck/England/041295/2022_H5N1
A/domestic_duck/England/105414/2022_H5N1
A/domestic_duck/England/104859/2022_H5N1
A/chicken/England/046274/2022_H5N1
A/Fox/Netherlands/EMC1/2022_H5N1
A/Barnacle_goose/Netherlands/2/2022_H5N1
A/Fox/Netherlands/EMC3/2022_H5N1
A/black-backed_gull/Netherlands/22006711-001/2022_H5N1
A/Sandwich_Tern/Netherlands/6/2022_H5N1
A/Common_Tern/Netherlands/7/2022_H5N1
A/Sandwich_Tern/Netherlands/2/2022_H5N1
A/sandwich_tern/Netherlands/22010437-002/2022_H5N1
A/Caspian_Gull/Netherlands/2/2022_H5N1
A/Barnacle_goose/Netherlands/5/2022_H5N1
A/goose/Netherlands/22000487-001/2022_H5N1
0.1

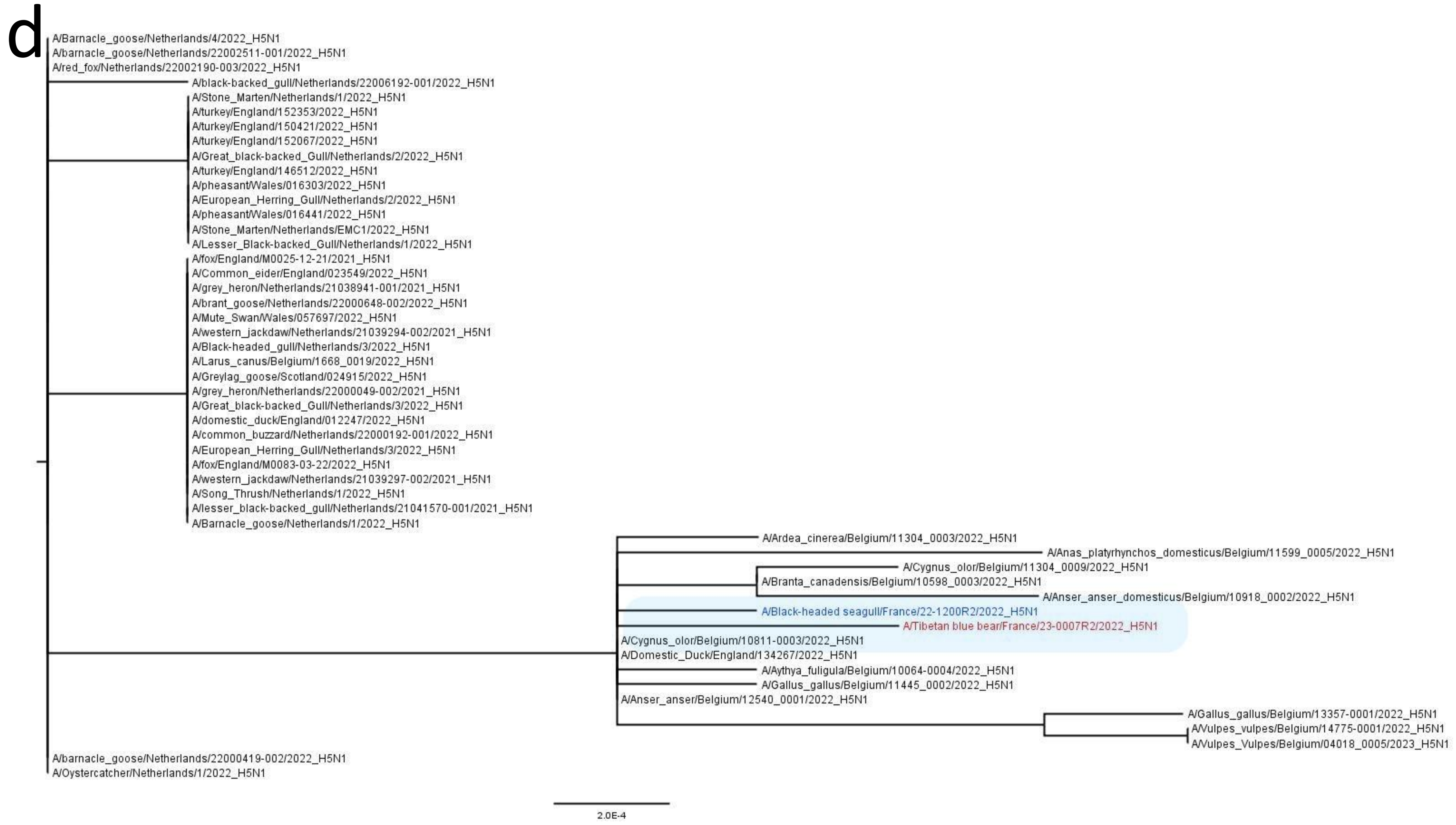
d
A/Barnacle_goose/Netherlands/4/2022_H5N1
A/barnacle_goose/Netherlands/22002511-001/2022_H5N1
A/red_fox/Netherlands/22002190-003/2022_H5N1
A/black-backed_gull/Netherlands/22006192-001/2022_H5N1
A/Stone_Marten/Netherlands/1/2022_H5N1
A/turkey/England/152353/2022_H5N1
A/turkey/England/150421/2022_H5N1
A/turkey/England/152067/2022_H5N1
A/Great_black-backed_Gull/Netherlands/2/2022_H5N1
A/turkey/England/146512/2022_H5N1
A/pheasant/Wales/016303/2022_H5N1
A/European_Herring_Gull/Netherlands/2/2022_H5N1
A/pheasant/Wales/016441/2022_H5N1
A/Stone_Marten/Netherlands/EMC1/2022_H5N1
A/Lesser_Black-backed_Gull/Netherlands/1/2022_H5N1
A/fox/England/M0025-12-21/2021_H5N1
A/Common_eider/England/023549/2022_H5N1
A/grey_heron/Netherlands/21038941-001/2021_H5N1
A/brant_goose/Netherlands/22000648-002/2022_H5N1
A/Mute_Swan/Wales/057697/2022_H5N1
A/western_jackdaw/Netherlands/21039294-002/2021_H5N1
A/Black-headed_gull/Netherlands/3/2022_H5N1
A/Larus_canus/Belgium/1668_0019/2022_H5N1
A/Greylag_goose/Scotland/024915/2022_H5N1
A/grey_heron/Netherlands/22000049-002/2021_H5N1
A/Great_black-backed_Gull/Netherlands/3/2022_H5N1
A/domestic_duck/England/012247/2022_H5N1
A/common_buzzard/Netherlands/22000192-001/2022_H5N1
A/European_Herring_Gull/Netherlands/3/2022_H5N1
A/fox/England/M0083-03-22/2022_H5N1
A/western_jackdaw/Netherlands/21039297-002/2021_H5N1
A/Song_Thrush/Netherlands/1/2022_H5N1
A/lesser_black-backed_gull/Netherlands/21041570-001/2021_H5N1
A/Barnacle_goose/Netherlands/1/2022_H5N1
A/Ardea_cinerea/Belgium/11304_0003/2022_H5N1
A/Anas_platyrhynchos_domesticus/Belgium/11599_0005/2022_H5N1
A/Cygnus_olor/Belgium/11304_0009/2022_H5N1
A/Branta_canadensis/Belgium/10598_0003/2022_H5N1
A/Anser_anser_domesticus/Belgium/10918_0002/2022_H5N1
A/Black-headed seagull/France/22-1200R2/2022_H5N1
A/Tibetan blue bear/France/23-0007R2/2022_H5N1
A/Cygnus_olor/Belgium/10811-0003/2022_H5N1
A/Domestic_Duck/England/134267/2022_H5N1
A/Aythya_fuligula/Belgium/10064-0004/2022_H5N1
A/Gallus_gallus/Belgium/11445_0002/2022_H5N1
A/Anser_anser/Belgium/12540_0001/2022_H5N1
A/Gallus_gallus/Belgium/13357-0001/2022_H5N1
A/Vulpes_vulpes/Belgium/14775-0001/2022_H5N1
A/Vulpes_Vulpes/Belgium/04018_0005/2023_H5N1
A/barnacle_goose/Netherlands/22000419-002/2022_H5N1
A/Oystercatcher/Netherlands/1/2022_H5N1
2.0E-4

e

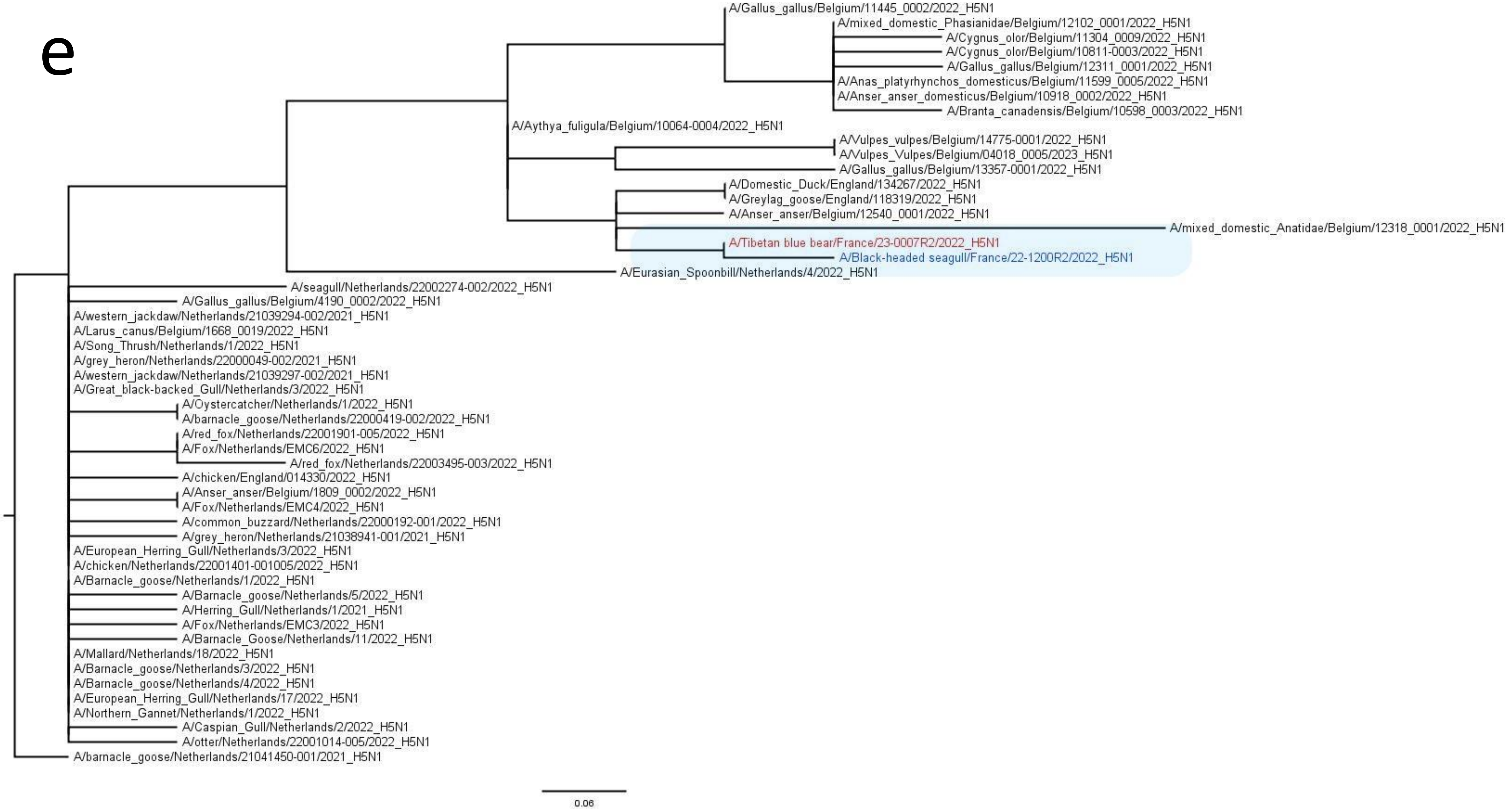
A/Gallus_gallus/Belgium/11445_0002/2022_H5N1
A/mixed_domestic_Phasianidae/Belgium/12102_0001/2022_H5N1
A/Cygnus_olor/Belgium/11304_0009/2022_H5N1
A/Cygnus_olor/Belgium/10811-0003/2022_H5N1
A/Gallus_gallus/Belgium/12311_0001/2022_H5N1
A/Anas_platyrhynchos_domesticus/Belgium/11599_0005/2022_H5N1
A/Anser_anser_domesticus/Belgium/10918_0002/2022_H5N1
A/Branta_canadensis/Belgium/10598_0003/2022_H5N1
A/Aythya_fuligula/Belgium/10064-0004/2022_H5N1
A/Vulpes_vulpes/Belgium/14775-0001/2022_H5N1
A/Vulpes_Vulpes/Belgium/04018_0005/2023_H5N1
A/Gallus_gallus/Belgium/13357-0001/2022_H5N1
A/Domestic_Duck/England/134267/2022_H5N1
A/Greylag_goose/England/118319/2022_H5N1
A/Anser_anser/Belgium/12540_0001/2022_H5N1
A/mixed_domestic_Anatidae/Belgium/12318_0001/2022_H5N1
A/Tibetan blue bear/France/23-0007R2/2022_H5N1
A/Black-headed seagull/France/22-1200R2/2022_H5N1
A/Eurasian_Spoonbill/Netherlands/4/2022_H5N1
A/seagull/Netherlands/22002274-002/2022_H5N1
A/Gallus_gallus/Belgium/4190_0002/2022_H5N1
A/western_jackdaw/Netherlands/21039294-002/2021_H5N1
A/Larus_canus/Belgium/1668_0019/2022_H5N1
A/Song_Thrush/Netherlands/1/2022_H5N1
A/grey_heron/Netherlands/22000049-002/2021_H5N1
A/western_jackdaw/Netherlands/21039297-002/2021_H5N1
A/Great_black-backed_Gull/Netherlands/3/2022_H5N1
A/Oystercatcher/Netherlands/1/2022_H5N1
A/barnacle_goose/Netherlands/22000419-002/2022_H5N1
A/red_fox/Netherlands/22001901-005/2022_H5N1
A/Fox/Netherlands/EMC6/2022_H5N1
A/red_fox/Netherlands/22003495-003/2022_H5N1
A/chicken/England/014330/2022_H5N1
A/Anser_anser/Belgium/1809_0002/2022_H5N1
A/Fox/Netherlands/EMC4/2022_H5N1
A/common_buzzard/Netherlands/22000192-001/2022_H5N1
A/grey_heron/Netherlands/21038941-001/2021_H5N1
A/European_Herring_Gull/Netherlands/3/2022_H5N1
A/chicken/Netherlands/22001401-001005/2022_H5N1
A/Barnacle_goose/Netherlands/1/2022_H5N1
A/Barnacle_goose/Netherlands/5/2022_H5N1
A/Herring_Gull/Netherlands/1/2021_H5N1
A/Fox/Netherlands/EMC3/2022_H5N1
A/Barnacle_Goose/Netherlands/11/2022_H5N1
A/Mallard/Netherlands/18/2022_H5N1
A/Barnacle_goose/Netherlands/3/2022_H5N1
A/Barnacle_goose/Netherlands/4/2022_H5N1
A/European_Herring_Gull/Netherlands/17/2022_H5N1
A/Northern_Gannet/Netherlands/1/2022_H5N1
A/Caspian_Gull/Netherlands/2/2022_H5N1
A/otter/Netherlands/22001014-005/2022_H5N1
A/barnacle_goose/Netherlands/21041450-001/2021_H5N1
0.06

f

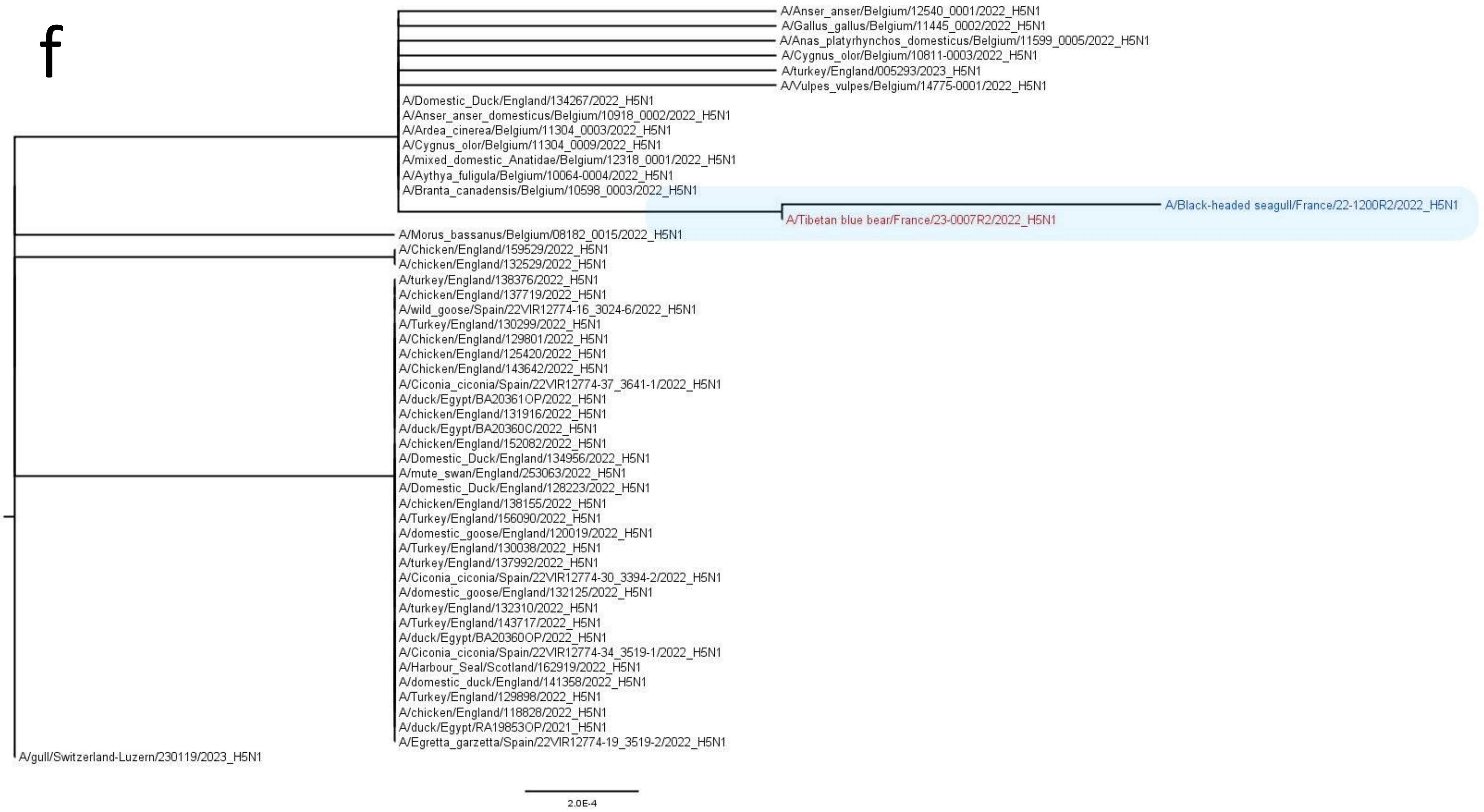
A/Anser_anser/Belgium/12540_0001/2022_H5N1
A/Gallus_gallus/Belgium/11445_0002/2022_H5N1
A/Anas_platyrhynchos_domesticus/Belgium/11599_0005/2022_H5N1
A/Cygnus_olor/Belgium/10811-0003/2022_H5N1
A/turkey/England/005293/2023_H5N1
A/Vulpes_vulpes/Belgium/14775-0001/2022_H5N1
A/Domestic_Duck/England/134267/2022_H5N1
A/Anser_anser_domesticus/Belgium/10918_0002/2022_H5N1
A/Ardea_cinerea/Belgium/11304_0003/2022_H5N1
A/Cygnus_olor/Belgium/11304_0009/2022_H5N1
A/mixed_domestic_Anatidae/Belgium/12318_0001/2022_H5N1
A/Aythya_fuligula/Belgium/10064-0004/2022_H5N1
A/Branta_canadensis/Belgium/10598_0003/2022_H5N1
A/Black-headed seagull/France/22-1200R2/2022_H5N1
A/Tibetan blue bear/France/23-0007R2/2022_H5N1
A/Morus_bassanus/Belgium/08182_0015/2022_H5N1
A/Chicken/England/159529/2022_H5N1
A/chicken/England/132529/2022_H5N1
A/turkey/England/138376/2022_H5N1
A/chicken/England/137719/2022_H5N1
A/wild_goose/Spain/22VIR12774-16_3024-6/2022_H5N1
A/Turkey/England/130299/2022_H5N1
A/Chicken/England/129801/2022_H5N1
A/chicken/England/125420/2022_H5N1
A/Chicken/England/143642/2022_H5N1
A/Ciconia_ciconia/Spain/22VIR12774-37_3641-1/2022_H5N1
A/duck/Egypt/BA20361OP/2022_H5N1
A/chicken/England/131916/2022_H5N1
A/duck/Egypt/BA20360C/2022_H5N1
A/chicken/England/152082/2022_H5N1
A/Domestic_Duck/England/134956/2022_H5N1
A/mute_swan/England/253063/2022_H5N1
A/Domestic_Duck/England/128223/2022_H5N1
A/chicken/England/138155/2022_H5N1
A/Turkey/England/156090/2022_H5N1
A/domestic_goose/England/120019/2022_H5N1
A/Turkey/England/130038/2022_H5N1
A/turkey/England/137992/2022_H5N1
A/Ciconia_ciconia/Spain/22VIR12774-30_3394-2/2022_H5N1
A/domestic_goose/England/132125/2022_H5N1
A/turkey/England/132310/2022_H5N1
A/Turkey/England/143717/2022_H5N1
A/duck/Egypt/BA20360OP/2022_H5N1
A/Ciconia_ciconia/Spain/22VIR12774-34_3519-1/2022_H5N1
A/Harbour_Seal/Scotland/162919/2022_H5N1
A/domestic_duck/England/141358/2022_H5N1
A/Turkey/England/129898/2022_H5N1
A/chicken/England/118828/2022_H5N1
A/duck/Egypt/RA19853OP/2021_H5N1
A/Egretta_garzetta/Spain/22VIR12774-19_3519-2/2022_H5N1
A/gull/Switzerland-Luzern/230119/2023_H5N1
2.0E-4

h

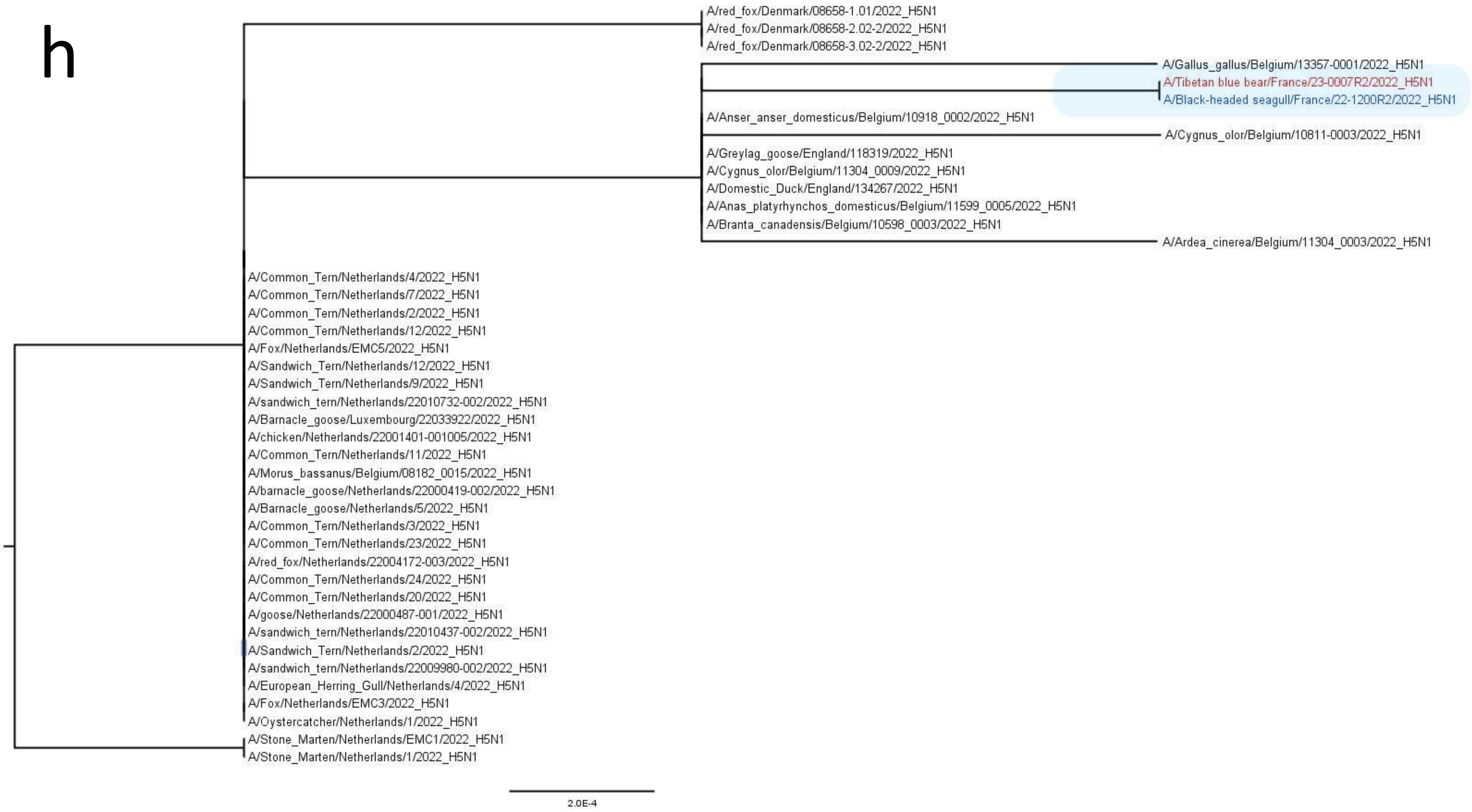
A/red_fox/Denmark/08658-1.01/2022_H5N1
A/red_fox/Denmark/08658-2.02-2/2022_H5N1
A/red_fox/Denmark/08658-3.02-2/2022_H5N1
A/Gallus_gallus/Belgium/13357-0001/2022_H5N1
A/Tibetan blue bear/France/23-0007R2/2022_H5N1
A/Black-headed seagull/France/22-1200R2/2022_H5N1
A/Anser_anser_domesticus/Belgium/10918_0002/2022_H5N1
A/Cygnus_olor/Belgium/10811-0003/2022_H5N1
A/Greylag_goose/England/118319/2022_H5N1
A/Cygnus_olor/Belgium/11304_0009/2022_H5N1
A/Domestic_Duck/England/134267/2022_H5N1
A/Anas_platyrhynchos_domesticus/Belgium/11599_0005/2022_H5N1
A/Branta_canadensis/Belgium/10598_0003/2022_H5N1
A/Ardea_cinerea/Belgium/11304_0003/2022_H5N1
A/Common_Tern/Netherlands/4/2022_H5N1
A/Common_Tern/Netherlands/7/2022_H5N1
A/Common_Tern/Netherlands/2/2022_H5N1
A/Common_Tern/Netherlands/12/2022_H5N1
A/Fox/Netherlands/EMC5/2022_H5N1
A/Sandwich_Tern/Netherlands/12/2022_H5N1
A/Sandwich_Tern/Netherlands/9/2022_H5N1
A/sandwich_tern/Netherlands/22010732-002/2022_H5N1
A/Barnacle_goose/Luxembourg/22033922/2022_H5N1
A/chicken/Netherlands/22001401-001005/2022_H5N1
A/Common_Tern/Netherlands/11/2022_H5N1
A/Morus_bassanus/Belgium/08182_0015/2022_H5N1
A/barnacle_goose/Netherlands/22000419-002/2022_H5N1
A/Barnacle_goose/Netherlands/5/2022_H5N1
A/Common_Tern/Netherlands/3/2022_H5N1
A/Common_Tern/Netherlands/23/2022_H5N1
A/red_fox/Netherlands/22004172-003/2022_H5N1
A/Common_Tern/Netherlands/24/2022_H5N1
A/Common_Tern/Netherlands/20/2022_H5N1
A/goose/Netherlands/22000487-001/2022_H5N1
A/sandwich_tern/Netherlands/22010437-002/2022_H5N1
A/Sandwich_Tern/Netherlands/2/2022_H5N1
A/sandwich_tern/Netherlands/22009980-002/2022_H5N1
A/European_Herring_Gull/Netherlands/4/2022_H5N1
A/Fox/Netherlands/EMC3/2022_H5N1
A/Oystercatcher/Netherlands/1/2022_H5N1
A/Stone_Marten/Netherlands/EMC1/2022_H5N1
A/Stone_Marten/Netherlands/1/2022_H5N1
2.0E-4
