## Supplementary Figures 1 to 3 for "High pathogenicity avian influenza A (H5N1) clade 2.3.4.4b virus infection in a captive Tibetan black bear (*Ursus thibetanus*): investigations based on paraffin-embedded tissues, France, 2022"

Additional illustrations of histopathological findings, viral antigen and RNA distribution in tissues from a tibetan blue bear and a black-headed seagull naturally-infected with H5N1 clade 2.3.4.4b HPAIV.

**Supplementary figure 1.** Additional illustrations of histopathological findings, viral antigen and RNA distribution in tissues from a Tibetan blue bear naturally-infected with H5N1 clade 2.3.4.4b HPAIV.

a. Heart: the myocardium appears within normal limits (H&E stain). b. Heart: there is mild non-specific background staining with no detection of viral antigen (IHC). c. Heart: sparse viral RNA detection is observed within the myocardial interstitium (insert), M gene RNAscope ISH. d. Spleen: thrombosis is observed within the splenic parenchyma (arrowhead) (H&E stain). e. Spleen: IHC reveals mild to moderate non-specific background staining with no significant detection of viral antigen. f. Spleen: rare detection of viral RNA is observed within the red pulp (M gene RNAscope ISH). g. Kidney: there is congestion and hemorrhages within the renal interstitium (H&E stain). h. Kidney: sparse viral antigen detection is observed within glomerular tufts (IHC). i. Kidney: no viral RNA is detected (RNAscope ISH). j. Liver: the hepatic parenchyma is multifocally replaced by nodular suppurative foci (arrowhead) (H&E stain). k. Liver: No viral antigen detection is observed (IHC). ). l. Liver: no viral RNA is detected (RNAscope ISH). Scale bars: 50µm (d-l) 100 µm, 200 µm (a-c).

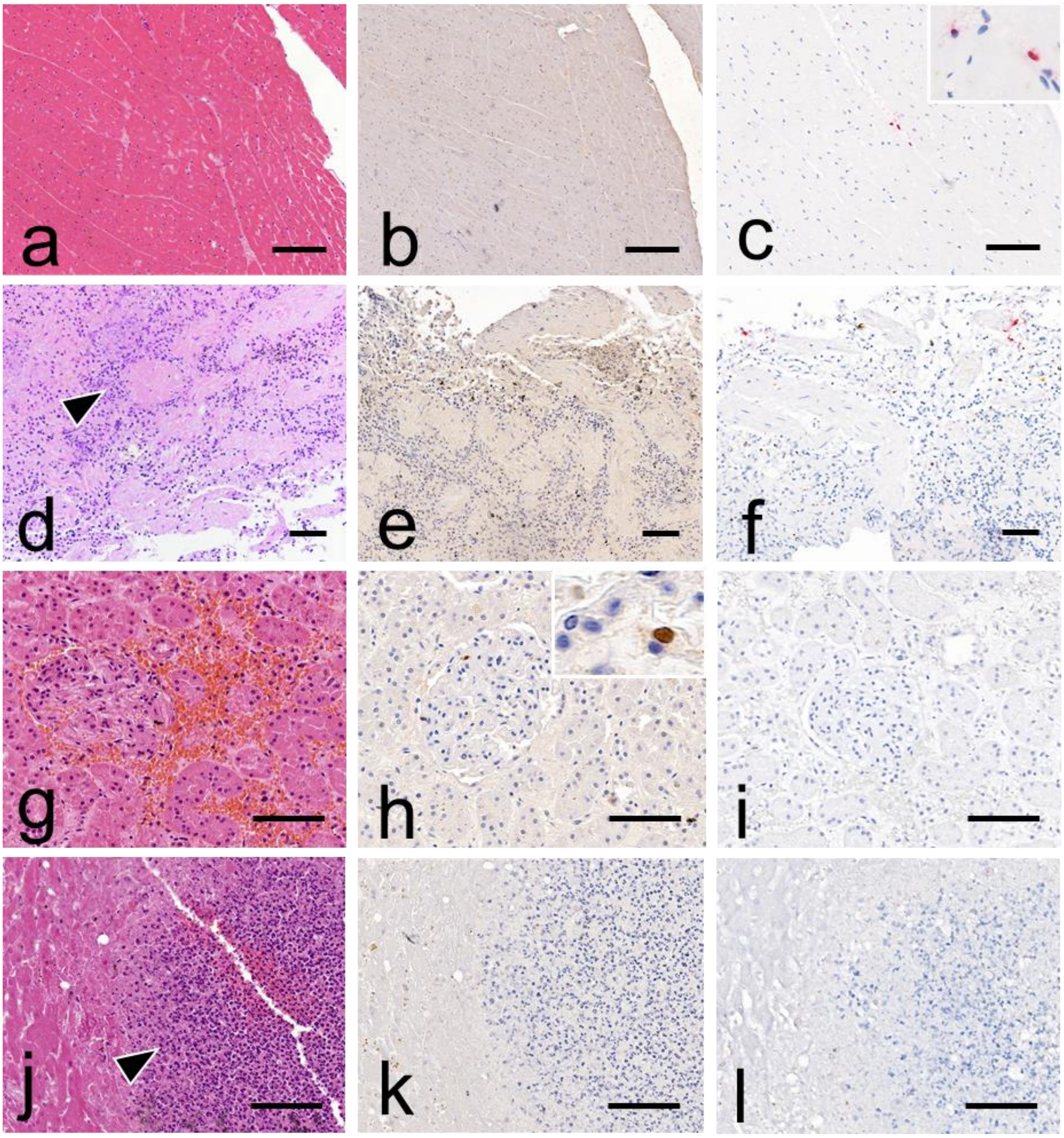

**Supplementary figure 2.** Additional illustrations of histopathological findings, viral antigenic and RNA distributions in tissues from a black-headed seagull naturally infected with H5N1 clade 2.3.4.4b HPAIV.

a. Optic Lobe (Central Nervous System, CNS): mild congestion is observed at a subgross view (H&E stain). b. Optic Lobe (CNS): extensive viral antigen detection involving neurons, neuropil, and glial cells (IHC). c. Optic Lobe (CNS): extensive viral RNA detection also involving neurons, glial cells and, to a lesser extent, neuropil (RNAscope ISH). d. Kidney: segmental multifocal epithelial necrosis involving renal tubules (arrowhead) (H&E stain). e. Kidney: multifocal segmental positive viral antigen detection within tubular nephrocytes (IHC). f. Kidney: multifocal segmental positive viral RNA detection within tubular nephrocytes (RNAscope ISH). g. Prerenal Nervous Ganglion: within normal limits (arrowhead) (H&E). h. Prerenal Nervous Ganglion: no viral antigen detection (IHC). i. Prerenal Nervous Ganglion: positive viral RNA detection within several neurons (RNAscope ISH). j. Intestine: the mucosa exhibits epithelial sloughing (arrowhead), while the outer layers appear within normal limits (H&E stain). k. Intestine: no viral antigen  is observed (IHC). l. Intestine: no viral RNA is detected (RNAscope ISH). Scale bars: 100 µm (d-l), 200 µm (a-c).

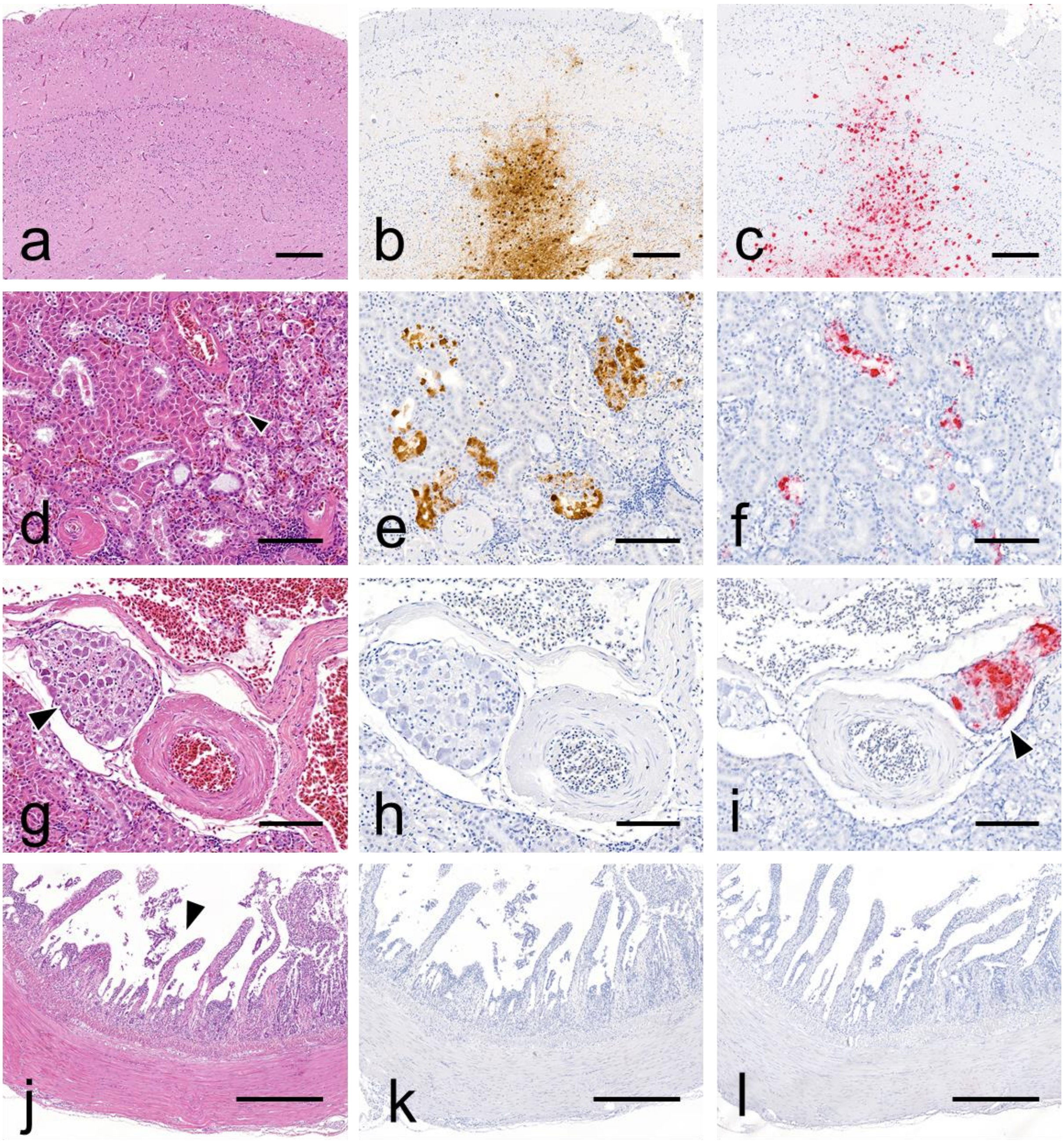

**Supplementary figure 3.** Additional illustrations of histopathological findings, viral antigen distribution in tissues from a black-headed seagull naturally-infected with H5N1 clade 2.3.4.4b HPAIV.

a. Thyroid Gland: degeneration and necrosis of thyroid follicles is observed (arrowhead) (H&E stain). b. Thyroid Gland: viral antigen detection involving both lining and sloughed epithelial cells of a single follicle (IHC). c. Heart: mild focal myocardial degeneration and necrosis (arrowhead) (H&E stain). d. Heart: multifocal viral antigen detection within cardiomyocytes (IHC). e. Lung: pulmonary lobules appear within normal limits (H&E stain). f. Lung: multifocal viral antigen detection within the capillary bed (IHC). g. Spleen and Pancreas (splenic Lobe): the splenic parenchyma (s) is within normal limits, while diffuse pancreatic necrosis is observed (p) (H&E stain). h. Spleen and Pancreas (Splenic Lobe): no viral antigen is observed in the spleen, while extensive positivity is present within the necrotic pancreatic parenchyma, presumably in exocrine acinar cells. Scale bars: 100 µm (a-f), 400 µm (g, h).

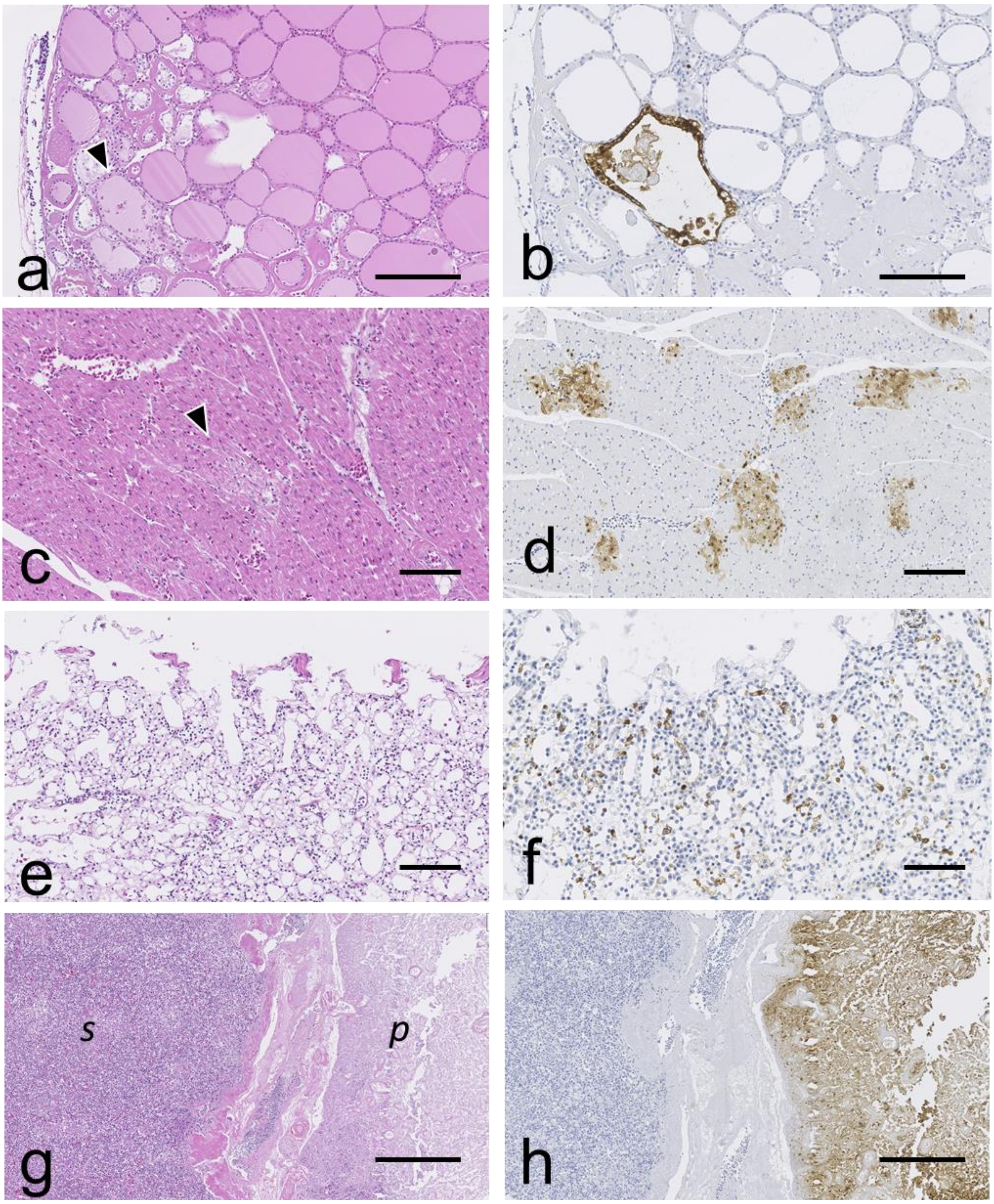
